# STARCH: Copy number and clone inference from spatial transcriptomics data

**DOI:** 10.1101/2020.07.13.188813

**Authors:** Rebecca Elyanow, Ron Zeira, Max Land, Benjamin J. Raphael

## Abstract

Tumors are highly heterogeneous, consisting of cell populations with both transcriptional and genetic diversity. These diverse cell populations are spatially organized within a tumor, creating a distinct tumor microenvironment. A new technology called *spatial transcriptomics* can measure spatial patterns of gene expression within a tissue by sequencing RNA transcripts from a grid of spots, each containing a small number of cells. In tumor cells, these gene expression patterns represent the combined contribution of regulatory mechanisms, which alter the rate at which a gene is transcribed, and genetic diversity, particularly copy number aberrations (CNAs) which alter the number of copies of a gene in the genome. CNAs are common in tumors and often promote cancer growth through upregulation of oncogenes or downregulation of tumor-suppressor genes. We introduce a new method STARCH (Spatial Transcriptomics Algorithm Reconstructing Copy-number Heterogeneity) to infer CNAs from spatial transcriptomics data. STARCH overcomes challenges in inferring CNAs from RNA-sequencing data by leveraging the observation that cells located nearby in a tumor are likely to share similar CNAs. We find that STARCH outperforms existing methods for inferring CNAs from RNA-sequencing data without incorporating spatial information.

## 1 Introduction

Intratumor heterogeneity, one of the hallmarks of cancer [Almendro et al., 2013], is characterized by distinct subpopulations of cells with both genetic and transcriptional diversity. These cell subpopulations, or *clones*, are spatially organized within a tumor. The clonal composition of a tumor may change over time or in response to therapy, and thus accurate quantification of intratumor heterogeneity is important for effective treatment of cancer. The genetic diversity within a tumor is due to somatic mutations that accumulate in the genome. A common type of somatic mutations – particularly in solid tumors – are copy number aberrations (*CNAs*) that amplify or delete contiguous regions in the genome. A sequence of CNAs is summarized in a copy number profile (*CNP*) which gives the number of copies of each region in the genome.

Numerous methods have been developed to infer CNPs from DNA sequencing data, with many of these methods building on early work deriving CNPs from microarrays [Olshen et al., 2004, Weir et al., 2007]. Methods to infer CNPs from next-generation DNA sequencing data examine differences between the observed and expected number of sequence reads that align to the genome. These methods leverage the observation that many CNAs are large, and thus the copy numbers of adjacent loci in the genome are often the same. Because of these positional dependencies, read counts from adjacent regions are correlated and thus the genome can be segmented into regions with the same copy number using change-point models [Xi et al., 2010] or a Hidden Markov Model (*HMM*) [Ha et al., 2012, 2014, Yu et al., 2017].

Inferring CNPs from RNA sequencing data is more challenging than from DNA sequencing data since the relative abundance of each transcript in a cell has many sources of variation beyond copy number [Tuch et al., 2010]. For this reason, CNP inference is typically not performed on RNA-seq data from bulk tumor samples due to the large number of samples that would be required to overcome this transcriptional variation. More recently, single cell RNA-sequencing (scRNA-seq) technologies enable measurement of mRNA transcripts in many cells from a single tumor, and several methods have been developed to infer CNPs from scRNA-seq data [Patel et al., 2014, Tirosh et al., 2016, Trinity-CTAT-Project, Fan et al., 2018]. In particular, InferCNV [Trinity-CTAT-Project] and HoneyBADGER [Fan et al., 2018] infer megabase or smaller CNAs from scRNA-seq data using 3-state Hidden Markov Models (HMMs) that mark the copy number of each gene as either a deletion, neutral, or an amplification. However, the inference of CNPs from scRNA-seq is complicated by the low sequencing coverage per cell, which limits the ability to distinguish variation in sequence coverage due to CNPs from other sources of transcriptional variation. A recent alternative approach uses both scRNA-seq and bulk DNA sequencing from the same tumor and assigns a CNP to each cell in the scRNA-seq data using a small number of copy number profiles derived from the bulk DNA sequencing data [Campbell et al., 2019, McCarthy et al., 2018, Andor et al., 2018]. These approaches rely on the availability of both types of sequencing data from the same tumor and also assume that the distinct samples from the same tumor largely share the same clonal organization, thus limiting their utility to specific datasets.

Recently *spatial transcriptomics* technologies have been developed to measure mRNA simultaneously across differential spatial locations, or spots, with each spot containing a small number (10-200) of cells [Ståhl et al., 2016]. Unlike bulk or single-cell RNA sequencing, spatial transcriptomics preserves the spatial location of each gene expression measurement, facilitating analysis of spatial patterns of gene expression. Spatial transcriptomics has been used to identify spatial patterns of gene expression in tissues [Ståhl et al., 2016, Liu et al., 2019], spatially distributed differentially expressed genes [Svensson et al., 2018, Arnol et al., 2019, Berglund et al., 2018] and spatially distributed cell clusters [Pettit et al., 2014, Zhu et al., 2018]. Here we propose a different use of the spatial information from spatial transcriptomics data: to identify CNPs and the spatial distribution of clones within a tumor sample.

We introduce a new method STARCH to jointly infer the CNPs and spatial locations of tumor clones using spatial transcriptomics data from a tumor. Similar to existing methods that infer CNPs from scRNA-seq, STARCH relies on the idea that most large CNAs result in correlated changes in gene expression across the multiple adjacent genes whose copy number is altered by the CNA. However, STARCH also leverages the observations that the CNAs in a tumor are related by a shared evolutionary history, and that nearby spots are likely to be genetically similar. These spatial correlations amplify the weak signal of CNPs in individual spots. STARCH models the spatial dependencies between clones in a tumor using a Hidden Markov Random Field (*HMRF*) and the positional correlations between copy numbers of adjacent genes using an HMM. We show on simulated tumor spatial transcriptomics data that STARCH outperforms existing approaches for CNA inference from transcriptomics data. On spatial transcriptomics data from four adjacent layers of a breast tumor, STARCH derives clones and CNPs that are more coherent across layers compared to existing methods.

## 2 Methods

### 2.1 Problem Definition

To identify CNAs in spatial transcriptomics data, we rely on several assumptions about the data and the underlying biological process: (1) The sequenced sample contains a small number of tumor clones and each clone is characterized by a distinct copy number profile; (2) CNAs that distinguish clones span multiple genes, creating dependencies between the copy numbers of adjacent genes; (3) Most of the cells in a spot belong to the same clone; (4) Nearby spots are likely to contain cells from the same clone, leading to spatial dependencies between the clone assignment for each spot.

These assumptions are well supported by biological evidence. First, tumors evolve through repeated clonal expansions [Nowell, 1976], and most tumors have a small number of subpopulations sharing similar sets of mutations and CNAs [Gerstung et al., 2020]. Second, the median size of CNAs in solid tumors is reported to be approximately 700Kb [Zack et al., 2013], which is significantly longer than the median size (24Kb) of a gene [Fuchs et al., 2014]. Third, several studies have demonstrated that nearby cells are likely to have the same somatic mutations [Merlo et al., 2006, Navin et al., 2010]. Thus, both the cells within each spot as well as the cells in nearby spots are likely to belong to the same clone.

We use these assumptions to process the spatial transcriptomics data for our model. First, based on the assumption that CNAs span many genes, we reduce much of the variance in expression attributed due to regulatory mechanisms by taking the average expression of multiple nearby genes into *bins* [Stranger et al., 2007] without losing information regarding their copy number status (Supplemental Section 2). Second, we scale the expression of each bin to be proportional to its copy number by normalizing the expression value by the expression value of the same bin in normal spots that do not harbor CNAs (Supplemental Section 1). We define **X** ∈ ℝ^*n*×*m*^ to be the resulting binned and normalized expression matrix from a spatial transcriptomics experiment, where *n* is the number of spots and *m* is the number of bins. Row **x**_*i*_ = [*x*_*i*1_,…,*x*_*im*_] represents the expression profile of spot *i*. We represent spatial dependencies between spots by a spot network *G* = (*V,E*), where *V* = {*v*_1_, …, *v*_*n*_} is the set of spots and an edge (*v*_*i*_,*v* _*j*_) ∈ *E* represents two neighboring spots (Figure 1(b)).

**Figure 1:**
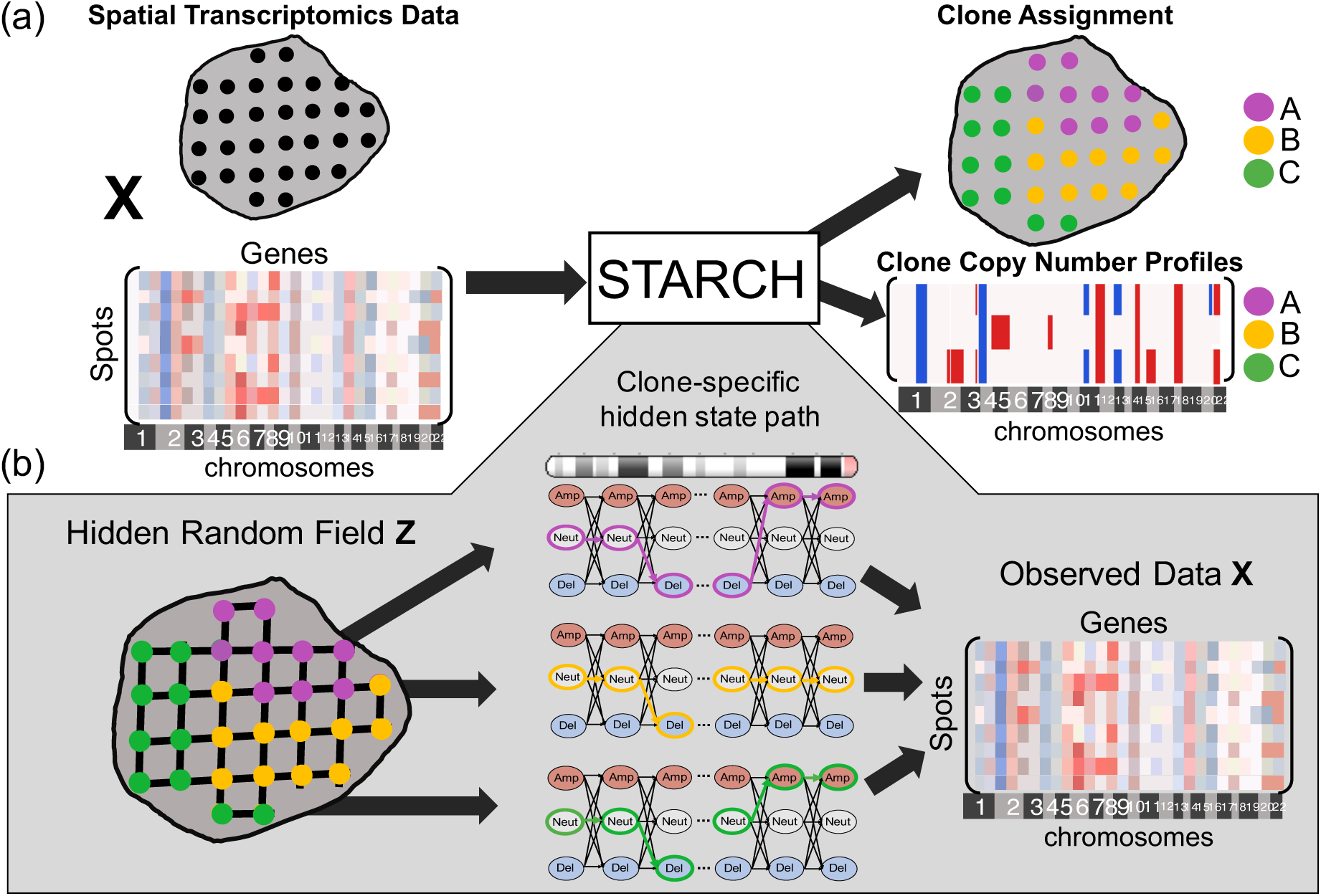
(a) **Overview of STARCH**. The inputs to STARCH are a spatial transcriptomics dataset consisting of a spot × gene expression matrix **X** and the spatial locations of each spot. The outputs are an assignment of each spot to one *K* clones and a copy number profile (CNP) for each clone. (b) **The STARCH model**. Clone assignments **Z** are modeled by a Hidden Markov Random Field (HMRF), where the hidden states are the clone assignments of the spots and the observed values are the transcript counts in **X**. The emission probabilities of each spot are computed from a Hidden Markov Model (HMM) that describes the CNP of each clone and chromosome arm.

Given a normalized expression matrix **X** ∈ ℝ^*n*×*m*^, a spot network *G* = (*V,E*), and a number *K* of clones, our goals are to: (1) infer a clone assignment matrix **Z** ∈ {0,1}^*n*×*K*^, where *z*_*ik*_ = 1 if spot *i* belongs to clone *k* and *z*_*ik*_ = 0 otherwise; (2) infer a copy number profile matrix **C** ∈ *S*^*K*×*m*^, where *S* is a set of possible copy number states and each row **c**_*k*_ = [*c*_*k*1_,…,*c*_*km*_] is the copy number profile of clone *k* (Figure 1).

### 2.2 STARCH model

STARCH aims to find a clone assignment matrix 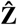 and a copy number profile matrix **Ĉ** that maximize the joint probability of the observed spot×bin expression matrix **X**,

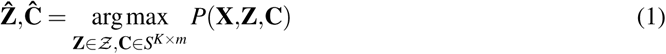

where 𝒵 is the set of all possible clone assignment matrices and *S*^*K*×*m*^ is the set of clone copy number profile matrices. We model the spatial dependencies between neighboring spots using a hidden Markov random field (HMRF), where the clone assignment **z**_*i*_ of a spot *i* depends only on the clone assignment of its neighboring spots. We model the dependencies between the copy numbers of adjacent bins along each chromosome arm using a hidden Markov model (HMM). The details of the HMRF and HMM are described in Sections 2.2.1 and 2.2.2 below.

#### 2.2.1 Hidden Markov Random Field (HMRF) for clone assignment matrix Z

We compute the joint probability *P*(**X,C,Z**) by conditioning on the clone assignment matrix **Z**,

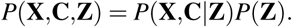

We model the clone assignment matrix **Z** as a configuration of a Markov random field on the spot network *G* = (*V,E*). Each row **z**_*i*_ denotes the clone assignment of spot *i*, where *z*_*ik*_ = 1 if spot *i* is from clone *k*, and *z*_*ik*_ = 0 otherwise. Note that **z**_*i*_ has exactly one entry equal to 1. Let *N*_*i*_ = {*v*_*j*_|(*v*_*i*_,*v* _*j*_) ∈ *E* be the set of neighbors of node *v*_*i*_. We assume that the clone assignment **z**_*i*_ of vertex *v*_*i*_ depends only on the clone assignments **z** _*j*_ of its neighboring vertices *v* _*j*_ ∈ *N*_*i*_. Thus, **Z** obeys the local Markov property:

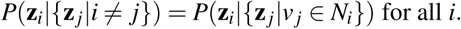

We model the observed gene expression matrix **X** and the clone CNP matrix **C** by a Hidden Markov Random Field (HMRF) [Zhang et al., 2001] whose hidden states are given by the clone assignment matrix **Z**. An HMRF model is characterized by three properties: (1) the **unobserved random field** (**Z**) exhibits the local Markov property, (2) the random variables of the **observed random field** (**X,C**) are **conditionally independent** given a configuration of the hidden random field, and (3) the random variables in the observable random field follow a known emission probability distribution given a configuration of the hidden random field.

Following property (1), the probability *P*(**Z**) of the clone assignment configuration **Z** follows a Gibbs distribution [Kinderman and Snell, 1980]:

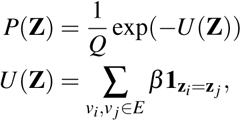

where *Q* is a normalizing constant, **1**_*t*_ is the indicator function, and *β* ≥ 0 is a parameter. Note that when *β* = 0, 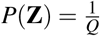 is constant and the assignment of each spot is independent of all other spots.

We assume that the CNP **c**_*k*_ of clone *k* depends only on spots assigned to clone *k* in the assignment matrix **Z**. Therefore, following property (2), the probability of the observed data **X** and the copy number profile matrix **C** given the hidden configuration **Z** is given by

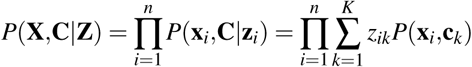

We compute *P*(**x**_*i*_, **c**_*k*_) using a Hidden Markov Model (HMM) as described in Section 2.2.2 below.

#### 2.2.2 Hidden Markov Model (HMM) for copy number profile matrix C

We compute the joint probability *P*(**x**_*i*_, **c**_*k*_) of gene expression **x**_*i*_ and copy number profile **c**_*k*_ for spot *i* using a Hidden Markov Model (HMM). We make two simplifying assumptions: (1) the average gene expression in bins from one chromosome arm is independent of the average gene expression in bins from other chromosome arms; (2) the copy number profile **c**_*k*_ of a clone *k* is independent of the copy number profiles of other clones. The first assumption follows from the observations that CNAs typically do not span across different chromosomes or chromosome arms. The second assumption simplifies the computation of the model and we find in practice is not limiting, although in reality there likely are dependencies between copy number profiles due to shared ancestry of the clones.

We model the expression profile **x**_*i*_ = [*x*_*i*1_,…,*x*_*im*_] of spot *i* on a chromosome arm as an ordered sequence of observed states from an HMM. The HMM contains three hidden copy number states *S* = {Deletion, Neutral, Amplification}. “Neutral” represents the diploid state (two copies), “Deletion” represents either zero or one copy, and “Amplification” represents any number of copies greater two. A three state HMM is also used in InferCNV [Trinity-CTAT-Project] and HoneyBADGER [Fan et al., 2018], two methods that infer CNPs from scRNA-seq data. The copy number states **c** follow a first-order Markov chain where the probability *P*(*c*_*j*+1_ = *b*|*c*_1_, …, *c*_*m*_) that bin *j* + 1 has copy number state *b* given the copy number states of all bins depends only on the copy number state at bin *j*; i.e., *P*(*c*_*j*+1_ = *b*|*c*_1_, …, *c*_*m*_) = *P*(*c*_*j*+1_ = *b*|*c* _*j*_ = *a*) = *t*_*ab*_. Let *T* = [*t*_*ab*_] be the 3 × 3 probability transition matrix, and let *π* = {*P*[*c*_1_ = *s*]|*s* ∈ *S*} denote the initial probability distribution. Finally, we define the emission probability *P*(*x*|*c* = *s*) of observing gene expression value *x* given copy number state *s* using a normal distribution with mean *µ*_*s*_ and standard deviation *σ*_*s*_. Thus, the joint probability *P*(**x**_*i*_,**c**) is given by

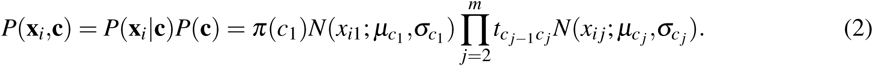

We denote by Θ = ∪_*s*∈*S*_{*µ*_*s*_,*σ*_*s*_} the set of parameters for the emission probability. Thus, the set *λ* = {*T,π*,Θ} of parameters fully specifies the HMM.

### 2.3 STARCH algorithm

Since it is computationally infeasible to maximize the posterior probability (Eq. 1) over all possible clone assignments and copy number profile matrices, we use an alternating coordinate ascent approach that iteratively updates **C** conditioned on **Z**, and vice-versa:

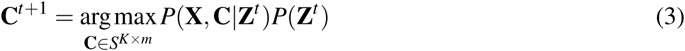

and

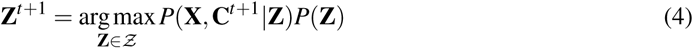

where **C**^*t*^ and **Z**^*t*^ are estimates of the clone copy number and assignment matrices for iteration *t*. Notice that since **Z**^*t*^ is fixed in iteration *t* + 1, the probability *P*(**Z**^*t*^) is constant and does not affect the maximization over **C**. Prior to performing coordinate ascent, we learn the values of the HMM parameters *λ* using a generalized version of the Baum-Welch algorithm [Li et al., 2000] and the expression profiles **x**_*I*_ of all spots.

In each iteration of the coordinate ascent, we compute **C**^*t*+1^ and **Z**^*t*+1^ as follows. First, we compute **C**^*t*+1^ given the clone assignments **Z**^*t*^ (equation (3)) using the Viterbi algorithm. Specifically, for each clone and chromosome arm we compute the state path **c**_*k*_ in the HMM that maximizes the joint probability of the observed expression profiles **x**_*i*_ of all spots *i* that are assigned to clone *k*. We compute **Z**^*t*+1^ using the Iterated Conditional Modes (ICM) algorithm [Besag, 1974]. ICM finds a local maximum of Equation 4 by iteratively updating each 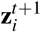 conditioned on the clone assignment 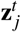 of each neighbor *j* of *i* from the previous iteration. We alternate between computing **C**^*t*+1^ given **Z**^*t*^ and **Z**^*t*+1^ given **C**^*t*^ until convergence or a maximum number of iterations is reached (see Supplemental Section 3).

## 3 Results

We evaluated STARCH by analyzing three synthetic spatial transcriptomics datasets and a spatial transcriptomics dataset from four sections of a breast tumor [Ståhl et al., 2016]. On each dataset, we compare STARCH (*β* = 2) to a version of STARCH that does not use spatial information (*β* = 0), and to InferCNV [Trinity-CTAT-Project], one of the standard methods for deriving CNAs from scRNA-seq data. InferCNV infers copy number profiles using a 3-state HMM and infers the clone assignments by performing hierarchical clustering on the expression profiles.

### 3.1 Synthetic spatial transcriptomics data

We compare STARCH and InferCNV on three synthetic spatial transcriptomics datasets. To ensure that tumor clones and their associated copy number profiles are biologically reasonable, we use tumor clones and copy number profiles derived from paired single-cell DNA sequencing and single-cell RNA-sequencing data of breast and ovarian tumors from Campbell et al. [2019].

First, we create a synthetic spatial transcriptomics dataset containing 150 spots with clones and copy number profiles derived from the published analysis of a triple-negative breast cancer patient-derived xenograft (SA501) from Campbell et al. [2019]. We assume that the gene expression at each spot is derived from one of 3 clones (labeled `A`, `B`, and `C`) whose copy number profiles are obtained from the published copy numbers. Specifically, we assign each genomic region with copy number less than 2 to be a deletion and each genomic region with copy number greater than 2 to be an amplification. We arrange 150 spots in a 12 × 13 grid, and assign contiguous block of spots to the same clone to simulate a spatial distribution of clones (Figure 2(a)). We generate a 150 × 6557 expression matrix **X** = [*x*_*ij*_] where each entry 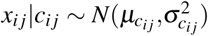 is normally distributed according to its copy number state *c*_*ij*_ ∈ {Deletion,Neutral,Amplification}. For each copy number state *c* ∈ {Deletion,Neutral,Amplification}, we estimate values of the mean *µ*_*c*_ and variance 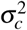 from the scRNA-seq data from the same triple-negative breast cancer patient-derived xenograft from [Campbell et al., 2019]. We select 25 of the 75 spots from clone `A` to represent normal spots with a diploid copy number profile, and use these spots to normalize the simulated expression matrix **X**, which contains a total of 125 spots.

**Figure 2:**
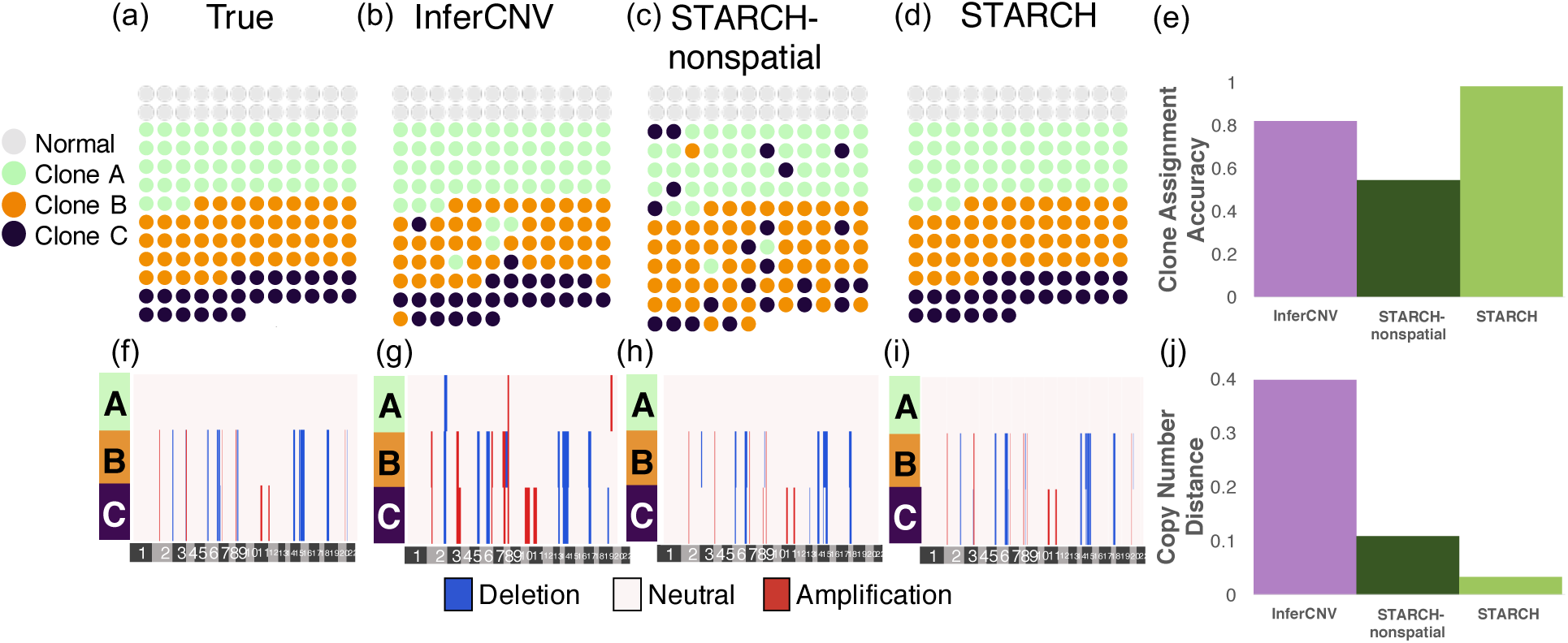
Clone assignment and CNA inference results from InferCNV, STARCH-nonspatial, and STARCH on synthetic spatial transcriptomics data. (a) Simulated spatial structure for clones `A` (green), `B` (orange) and `C` (purple). Normal spots are indicated in gray. (b-d) Clone assignments inferred by each method. (e) Clone assignment accuracy for each method computed as the Adjusted Rand Index between true and inferred clone assignments (higher is better) (f-i). Inferred copy number profiles for each clone. (j) Copy number distance between true and inferred copy number profiles (lower is better).

We ran STARCH and InferCNV using the matrix **X** and with *K* = 3 clones. We ran STARCH both with the spatial information (*β* = 2) and without (*β* = 0, denoted *STARCH-nonspatial*). We evaluate the performance of each method using two measures: (1) the *clone assignment accuracy* computed as the adjusted rand index (ARI) between true and inferred clone assignments; (2) the *copy number distance* between true and inferred copy number profiles using the average normalized Hamming distance (Supplemental Section 4). We find that STARCH outperforms both STARCH-nonspatial and InferCNV in assigning spots to their respective clones (clone assignment accuracy of .98, .59, and .82 respectively) and recovering the simulated CNAs (copy number distance of .03,.12, and .39 respectively) (Figure 2(e)). InferCNV tends to overestimate the number of CNAs, identifying several CNAs in the diploid clone `A` (Figure 2(b)). In contrast, STARCH-nonspatial tends to underestimate the number of CNAs; for example STARCH-nonspatial identifies a deletion on chromosome 2 in clone `B` (Figure 2(c)) only while STARCH correctly identifies the deletion in both clones `B` and `C` (Figure 2(d)).

### 3.2 Pseudo-spatial transcriptomics from matched scDNA-seq and scRNA-seq

Next, we compare methods on two *pseudo-spatial transcriptomics* datasets that we generate by adding spatial structure to scRNA-seq data from breast and ovarian tumors whose clonal composition was derived from matched scDNA-seq from the same tumors [Campbell et al., 2019]. In contrast to the fully synthetic data above, here the expression matrix **X** comes directly from scRNA-seq data. Thus, CNA inference from this pseudo-spatial transcriptomics data is more difficult than the fully synthetic data because real scRNA-seq data contains additional variation in transcript counts due to true biological variation in gene expression between cells and dropout of lowly expressed genes due to sampling and other technical artifacts.

We first generate a pseudo-spatial transcriptomics dataset from matched scRNA-seq and DNA-seq data from a breast cancer patient-derived xenograft (SA501) from Campbell et al. [2019]. As in the synthetic dataset generated above, we use the three clones (`A`, `B`, and `C`) and their corresponding copy number profiles derived from scDNA-seq. We randomly select 200 of the 930 cells assigned to clone `A` and define them as normal cells which we use to normalize **X** to a diploid copy number profile. We assign 192 cells to clone `B`, and only 30 cells to clone `C`. The unequal clone proportions complicate CNA inference in this dataset. We assign each of the 1152 cells to a position on a 35 × 34 grid with two contiguous rectangular regions containing cells from clone `B`, one rectangular region containing cells from clone `C`, one region containing normal cells, and the remaining spots containing cells from clone `A` (Figure 3(a)).

**Figure 3:**
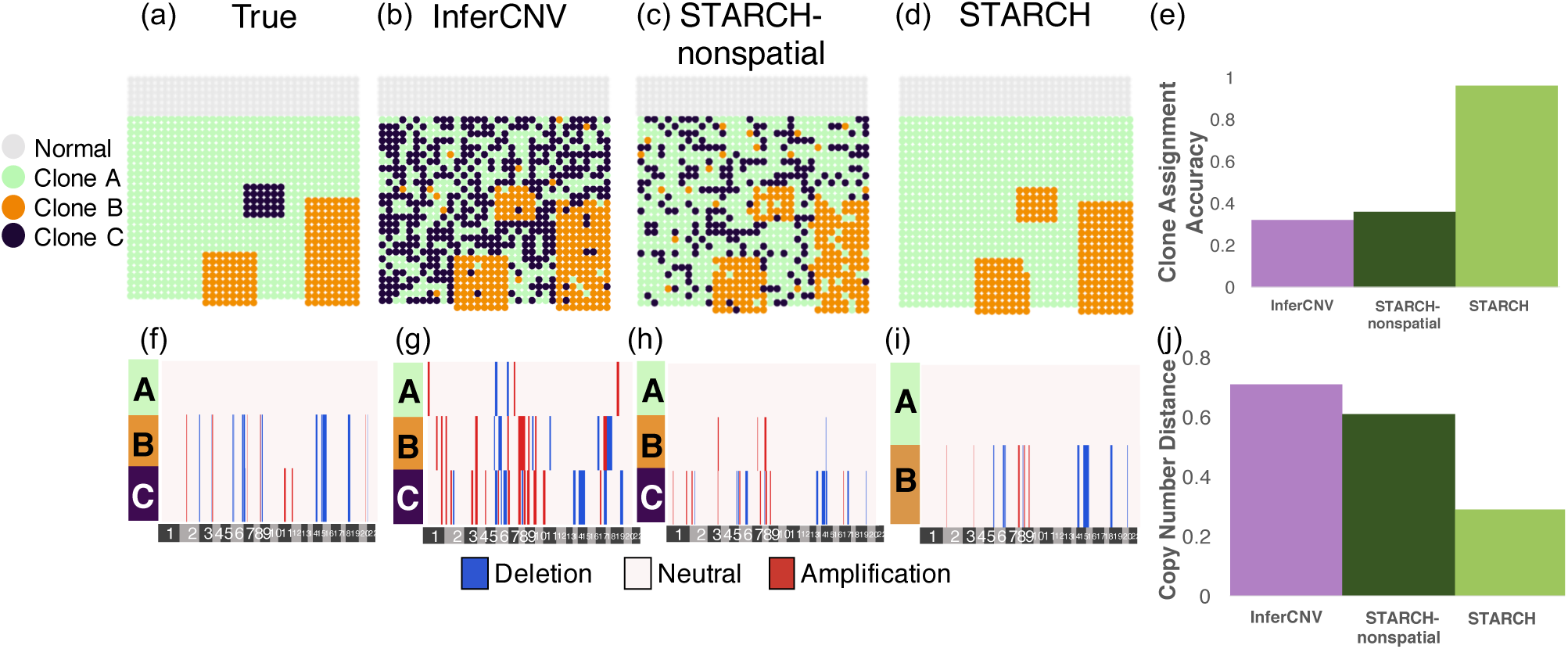
Clone assignment and CNA inference results from InferCNV, STARCH-nonspatial, and STARCH on pseudo-spatial data from patient-derived xenograft SA501 [Campbell et al., 2019]. (a) Simulated spatial structure for clones `A` (green), `B` (orange) and `C` (purple). Normal spots are indicated in gray. (b-d) Clone assignments inferred by each method. (e) Clone assignment accuracy for each method computed as the Adjusted Rand Index between true and inferred clone assignments (higher is better). (f-i) Inferred copy number profiles for each clone. (j) Copy number distance between true and inferred copy number profiles (lower is better).

We ran STARCH, STARCH-nonspatial, and InferCNV on the psuedo-spatial transcriptomics dataset using *K* = 3 clones and compared the copy number distance between true and inferred copy number profiles as well as the clone assignment accuracy between true and inferred clone assignments. We find that by incorporating spatial information STARCH achieved higher clone assignment accuracy and lower copy number distance than STARCH-nonspatial and InferCNV (Figure 3). STARCH infers the copy number profile of clones `A` and `B` with low error and assigns most of the spots to their respective clones (ARI .97), while InferCNV and STARCH-nonspatial identify many CNAs not present in the data and fail to correctly assign spots to clones (ARI of .32 and .46 respectively). We note that none of the methods are able to identify the 30 spots from clone `C`. STARCH assigned all of these 30 spots to clone `B`, which has a very similar copy number profile to clone `C` (differing from clone `B` by only two amplifications on chromosome 11). InferCNV and STARCH-nonspatial, however, assign these spots to three different clones, none of which match the true copy number profile of clone `C` (Figure 3). Furthermore, STARCH takes significantly less time to run than InferCNV, finishing in 11 minutes compared to 126 minutes for InferCNV using 4 cores (Supplemental Figure 2).

We derive a second pseudo-spatial transcriptomics dataset from matched scRNA-seq and DNA-seq data from a high grade serous carcinoma cell line OV2295 obtained from Campbell et al. [2019] (Supplemental section 8). This dataset contains two clones (`C` and `D`) as well as their corresponding copy number profiles derived from scDNA-seq. We performed the analogous comparison of methods on this pseudo-spatial transcriptomics dataset and found that by incorporating spatial information STARCH outperforms both STARCH-nonspatial and InferCNV in assigning spots to clones and inferring copy number profiles (Supplemental Figure 3).

### 3.3 Spatial transcriptomics data

Finally, we compare the performance of STARCH, STARCH-nonspatial and InferCNV on spatial transcriptomics data from Ståhl et al. [2016]. This dataset consists of spatial transcriptomics sequencing data from four adjacent layers of a breast cancer tissue biopsy.

We first ran STARCH on the spatial transcriptomics data from all four layers, with clones and copy number profiles shared across all four layers. Namely, we create a single gene expression matrix **X** whose rows are the spots from all four layers. Since the spatial coordinates are not aligned across layers, we define the spot network *G* as the union of disjoint grid networks for each layer; thus we model only spatial relationships within each layer. We developed a clustering method to differentiate tumor from normal spots in each layer (Supplemental section 9) and normalized the expression data relative to diploid normal spots identified by this method. The normal spots identified by this method (gray spots in Figure 4a) closely matched the regions labelled as non-cancerous on the imaging data from Ståhl et al. [2016].

**Figure 4:**
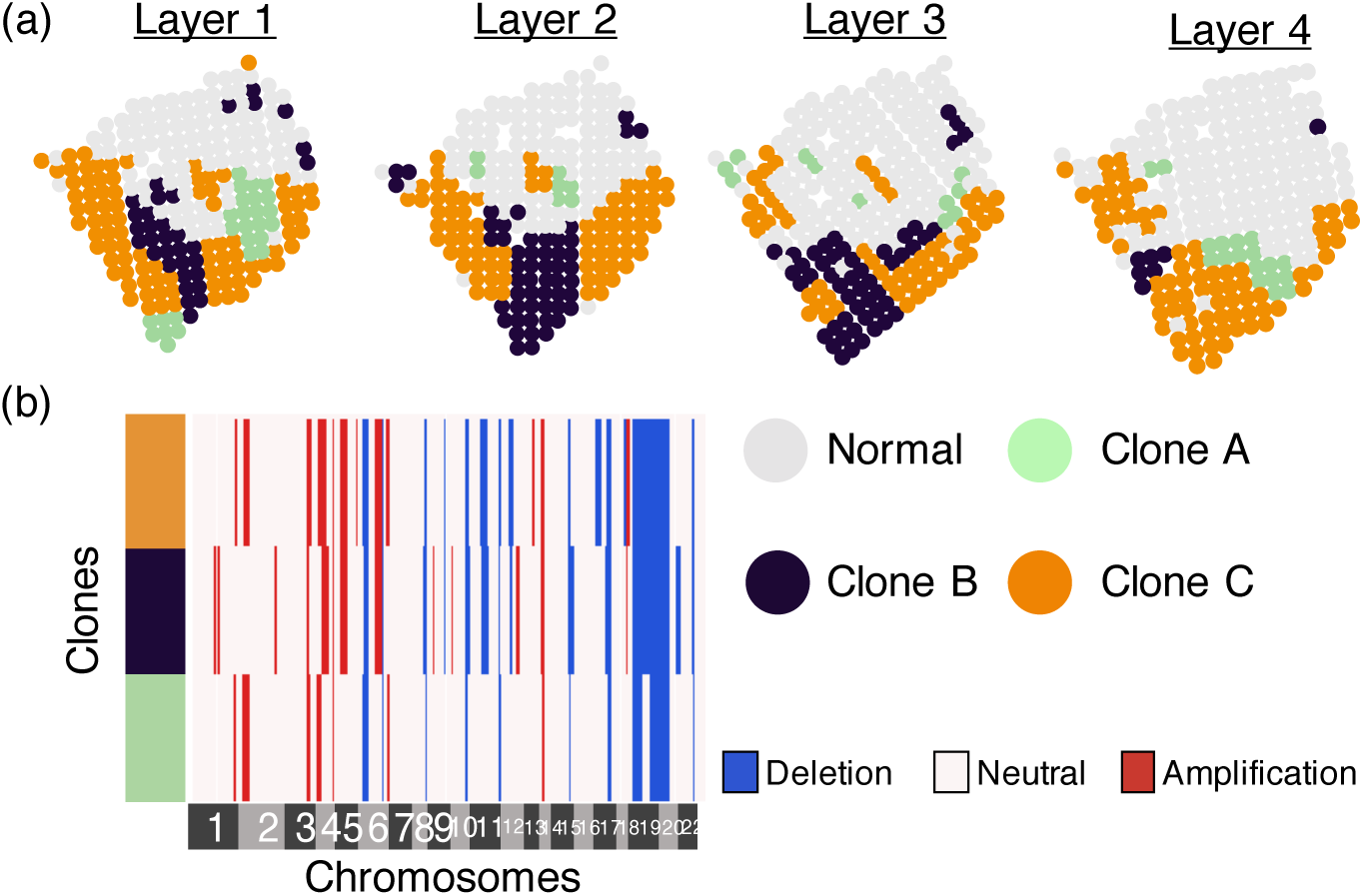
Clone assignments and copy number profiles inferred by running STARCH simultaneously on four layers of a breast cancer biopsy [Ståhl et al., 2016]. (a) Assignments of a normal clone (gray) and three tumor clones (green, blue, orange) for each layer. (b) Copy number profiles for each clone.

Running STARCH jointly across all four layers identifies three spatially distributed clones with distinct copy number profiles. Since we do not know the true clone assignments or copy number profiles, we cannot directly assess the accuracy of STARCH’s output in this dataset. However, STARCH’s results seems reasonable for two reasons. First, we observe that the clone assignments are visually aligned across adjacent layers, consistent with the expectation that genetically similarly clones should occupy contiguous spatial compartments [Sutherland, 1988]. Second, the copy number profiles of distinct clones share a large proportion of CNAs: 12 of the 35 distinct CNAs in the copy number profiles are clonal, i.e., present in all clones (Figure 4b). This number of clonal CNAs matches a study by Gao et al. [2016] who performed scDNA-seq of 12 breast cancer tumors and found 8 − 16 clonal CNAs per tumor. A large proportion of clonal CNAs are consistent with the punctuated evolution model, where most CNAs arise early in tumor evolution and prior to clonal diversification [Gao et al., 2016].

To more directly evaluate the performance of copy number inference methods on this data, we ran STARCH, STARCH-nonspatial, and InferCNV independently on the tumor spots from each layer. We compare the clone assignments and inferred copy number profiles between adjacent layers 1:2, 2:3, and 3:4 respectively. We find that the clone assignments inferred by STARCH are more consistent across adjacent layers of the tissue than the assignments inferred by the other two methods (Figure 5(a-c)). We also find that copy number profiles inferred by STARCH had lower average copy number distance between layers (.22,.17,.26) compared to InferCNV (.50,.33,.32) and STARCH-nonspatial (.38,.32,.30) (Figure 5(d-f)). These results are consistent with the expectation that the spatial locations of clones should be highly similar between adjacent layers. Furthermore, STARCH identifies 12 clonal CNAs conserved across all four layers, compared to only 9 such conserved clonal CNAs for STARCH-nonspatial and 2 for InferCNV. The presence of more clonal CNAs identified across all four layers is further evidence that STARCH has higher accuracy for CNA inference. Finally, STARCH identifies several subclonal CNAs present in a subpopulation of clones and observed across all four layers. One such subclonal CNA is a deletion on 12q13 in clones `B1`-`B4` that includes the tumor suppressor gene TP53. This deletion was identified across all four layers by STARCH, while the other two methods only identify this deletion in a subset of layers.

**Figure 5:**
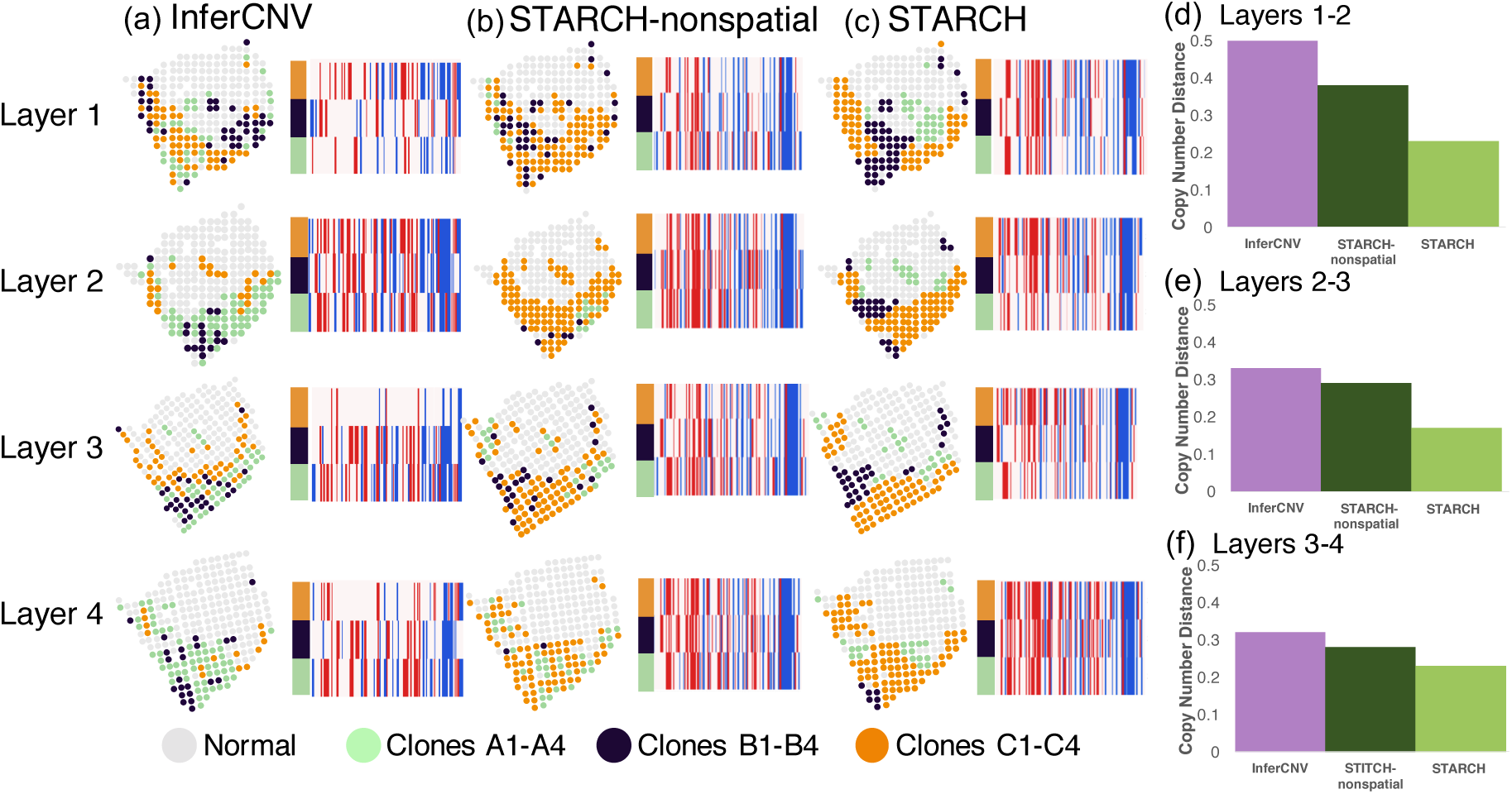
Clone assignment and CNA inference results from InferCNV, STARCH-nonspatial, and STARCH on spatial transcriptomics data from four layers of a breast cancer biopsy. For each method, CNA inference was performed separately for each of the four layers 1-4 ordered from top to bottom. (a-c) Clone assignment and copy number profiles from InferCNV, STARCH-nonspatial and STARCH. (d-f) Average copy number distance between layers 1 and 2, 2 and 3, and 3 and 4.

Finally, we investigate the performance of STARCH on a normal tissue where we do not expect to find copy number aberrations, providing a negative control. We ran STARCH on spatial transcriptomics of fresh frozen human heart tissue from a healthy donor obtained from 10X Genomics. When we ran STARCH with *K* = 3 clones, STARCH output a single clone containing only four small CNAs (Supplemental Figure 5), indicating a low false positive rate for CNA inference. Additionally, even though STARCH is given *K* = 3 clones as input, it converges to a single clonal population and does not infer erroneous subclones. These two results are promising and suggest that STARCH will produce few false positive results on real tumor spatial transcriptomics data. Full details of the analysis are in Supplemental Section 10.

## 4 Discussion

We propose a new method, STARCH, to infer CNAs from spatial transcriptomics data. STARCH jointly infers the assignment of cells to one of *K* clones and the CNPs of each clone by leveraging spatial dependencies between clone assignments for neighboring spots and regional dependencies between copy number across neighboring genes. To our knowledge, STARCH is the first method which utilizes spatial dependencies to infer clone assignment and CNPs. We demonstrate that by incorporating these spatial dependencies STARCH outperforms other non-spatial methods at recovering the true clone assignments and CNPs on simulated data. We also demonstrate that STARCH computes more plausible clone assignments and CNPs on real spatial transcriptomics data; STARCH’s clone assignments and CNPs are more consistent across adjacent layers of a breast cancer biopsy.

There are several limitations and areas for future improvement of STARCH. First, the copy number model used in STARCH is fairly coarse with only three copy number states (Deletion, Neutral, Amplification); a larger number of copy number states could provide more accurate CNPs and clone assignments. One interesting direction is to extend STARCH to infer allele-specific copy number, as has been done for bulk DNA-seq [Ha et al., 2014, Zaccaria and Raphael, 2018, Nik-Zainal et al., 2012], single-cell DNA-seq [Garvin et al., 2015, Zaccaria and Raphael, 2019], and single-cell RNA-seq [Fan et al., 2018]. Allele-specific copy number provides a more accurate representation of the clonal structure of tumors, is essential for identifying events such as copy-neutral loss of heterozygosity, and could be helpful in quantifying allele-specific gene expression. Second, the spatial model used in STARCH could be further refined. For example, rather than a 2D grid one might include additional edges such as diagonal edges or use weighted edges to represent distances between spots. For datasets with multiple layers, one could run STARCH with a 3D grid. In our current analysis, we did not connect any spots from different layers by an edge because the spatial coordinates for this dataset were not aligned in 3D. Finally, the copy numbers inferred by STARCH could be helpful to distinguish gene expression variation due to copy number aberrations from gene expression variation due to regulatory mechanisms, providing a more refined analysis of spatial transcriptomics data from cancer tissues containing numerous copy number aberrations.

## 5 Code Availability

STARCH is implemented in Python and is available at https://github.com/raphael-group/STARCH.

## 6 Acknowledgements

This work is supported by grant U24CA211000 from the US National Cancer Institute (NCI) and by grant 2018-182608 from the Chan Zuckerberg Initiative Donor-Advised Fund (DAF), an advised fund of Silicon Valley Community Foundation.

## Notes

### Competing Interest Statement

B.J.R. is a cofounder of, and consultant to, Medley Genomics

