## Supplemental Text for "STARCH: Copy number and clone inference from spatial transcriptomics data"

#### Contents

**1 STRNA-seq data processing** **2**

**2 Binning genes** **2**

**3 STARCH algorithm pseudocode** **2**

**4 Normalized Hamming Distance** **3**

**5 Selecting the value of parameter $\beta$ in the HMRF** **3**

**6 Model initialization** **3**

**7 Runtime** **4**

**8 Pseudo-spatial transcriptomics data from high grad serous carcinoma cell line OV2295** **4**

**9 Distinguishing cancer spots from normal spots** **5**

**10 Negative Control with human heart tissue** **7**

### 1 STRNA-seq data processing

We derive the input matrix $\mathbf{X}$ for STARCH from the raw STRNA-seq gene expression data using several processing steps, which aim to reduce the technical variability in the data, similar to InferCNV [Trinity-CTAT-Project]. First, genes with non-zero counts in fewer than 20 cells are removed. The data is then library-size normalized, where counts for each gene in a spot are divided by the sum of counts for the spot. This normalization removes variance due to differences in size or number of cells in each spot. We denote this normalized expression data by $\mathbf{G}$. Next, we log-normalized the data with pseudocount 1, computing $\log(\mathbf{G}+1)$. Then, we compute average expression values for genes in neighboring bins to reduce variation in expression from regulatory mechanisms acting at the gene level (see Section 2), while retaining large-scale variations due to copy number changes. We denote the resulting spot $\times$ bin expression matrix by $\mathbf{X}''$. To make the expression of each bin proportional to its copy number we subtract the median binned expression of normal spots from $\mathbf{X}''$. This step removes differences in total expression between bins and allow for direct comparison of expression across bins. We denote the resulting data matrix as $\mathbf{X}'$. Finally, the log-transformation is inverted giving our final normalized binned expression $\mathbf{X} = (\exp(\mathbf{X}') - 1)$.

### 2 Binning genes

To avoid high rates of false positive copy number calls due to the biological variability in transcript counts in STRNA-seq data, we average gene expression values across adjacent genes before running STARCH. Given a normalized gene expression matrix $\mathbf{G}$, we compute the average expression per bin, where each bin $b$ contains $w$ genes with a step size of $s$ genes. That is, for each bin $b$ and each spot $i$, the binned expression is

$$x''_{ib} = \frac{1}{w} \sum_{j=sb-w/2}^{j=sb+w/2} g_{ij},$$

where $g_{ij}$ is the normalized expression from spot $i$ that aligns to gene/transcript $j$. This results in a spot $\times$ bin expression matrix $\mathbf{X}''$. We bin based on genes rather than genomic intervals because the observed measurements are at the gene level and this allows measurements to be directly comparable across bins.

We choose a window size $w$ to balance the tradeoff between reducing expression variance due to technical noise and biological variability and identifying CNAs over reasonable size. The median length of a human gene is approximately 24Kb [Fuchs et al., 2014], with average intergenic distances between genes (excluding novel isoforms) being 14Kb or smaller [Djebali et al., 2012]. Comparatively, the median lengths for focal amplification and deletions have been reported to be approximately 900Kb and 700Kb respectively across many cancer types (with telomere-bounded focal CNAs being much longer at approximately 19.6Mb and 22.7Mb, respectively) [Zack et al., 2013]. Thus we expect the median CNA to span approximately 21 genes or more. For all experiments we use a window size $w$ such that there is a median of 5 transcripts per bin. This typically results in window size between 30 - 50 genes. For all experiments we use a step size of 1, so we get a copy number call for all $m$ genes in the dataset.

### 3 STARCH algorithm pseudocode

1. Process the raw data as described in Supplemental Section 1 to obtain the preprocessed spot $\times$ bin matrix $\mathbf{X}$.

2. Initialize parameters $\Theta$ and cluster assignments $\mathbf{Z}^0$ as described in Supplemental Section 6.

3. Learn the HMM parameters $\lambda$ using the Baum-Welch algorithm.

4. Estimate $\mathbf{C}^{t+1} = \arg\max_{\mathbf{C}\in S^{K\times m}} P(\mathbf{X},\mathbf{C}|\mathbf{Z}^t)P(\mathbf{Z}^t)$ using the Viterbi algorithm as described in Section 2.3

5. Estimate $\mathbf{Z}^{t+1} = \arg\max_{\mathbf{Z}\in\mathcal{Z}} P(\mathbf{X},\mathbf{C}^{t+1}|\mathbf{Z})P(\mathbf{Z})$ using the ICM algorithm as described in Section 2.3

6. Iterate between steps 3 and 4 until $\mathbf{C}^{t+1}$ converges or a maximum number of iterations (default 20) is reached.

### 4 Normalized Hamming Distance

To compare copy number profiles we define the *normalized Hamming distance* that gives more weight to differences in bins with non-neutral copy numbers, where a non-neutral bin is either an amplification or deletion. The Hamming distance $H(a,b)$ between two copy number profiles $a$ and $b$ is the total number of bins in which the two profiles differ. The Hamming distance between copy number profiles with few non-neutral bins will almost always be smaller than the Hamming distance between copy number profiles with many non-neutral bins – even if none of the bins match between profiles $a$ and $b$. To address this issue, we define the *normalized Hamming distance* to be

$$H_n(a,b) = \frac{H(a,b)}{|a|+|b|},$$

where $|a|$ and $|b|$ are the total number of non-neutral bins in copy number profiles $a$ and $b$ respectively. Thus, if all non-neutral bins are the same between $a$ and $b$, then $H_n(a,b) = 0$, and if all non-neutral bins differ between $a$ and $b$ then $H_n(a,b) = 1$.

### 5 Selecting the value of parameter $\beta$ in the HMRF

The parameter $\beta$ in the HMRF weighs the coherence between clone assignments of neighboring spots in the prior of the clone assignment matrix $\mathbf{Z}$. Higher values of $\beta$ give more spatially coherent cluster assignments, with neighboring spots more likely to be members of the same clone. Lower values of $\beta$ do not put much weight on the clone assignment of neighboring spots, but rather put more weight on fitting the clone assignment to the data (expression and copy number profiles).

We select the value of parameter $\beta$ used in our analysis using simulated data (see simulation setup in Section 3.1). In each simulation, we measured the distance between true and inferred CNA matrices $\mathbf{C}^*$ and $\hat{\mathbf{C}}$ as well as the difference between true and inferred clone assignment matrices $\mathbf{Z}^*$ and $\hat{\mathbf{Z}}$ over a range of $\beta$ values. We found that when we ran STARCH on the simulated data both the inferred CNAs and clone assignments were stable across a range of $\beta$ from 1 to 3 (Figure 1). The copy number distance between true and inferred CNA matrices, measured by normalized Hamming distance, as well as the difference between true and inferred clone assignment matrices, measured by the ARI, increased for $\beta$ values less than 1 or higher than 3 (Figure 1). We thus use $\beta = 2$ for all our analyses of simulated and real data.

### 6 Model initialization

In this section we describe how we obtain the initial clone assignment matrix $\mathbf{Z}^0$ and CNP matrix $\mathbf{C}^0$ as well as initial estimates of the HMM model parameters $\lambda$ which are then updated by the Baum-Welch algorithm. The clone assignment matrix $\mathbf{Z}^0$ is initialized by performing K-means clustering on the rows of the observed expression matrix $\mathbf{X}$. The number $K$ of clones may be selected either using prior knowledge or by computing

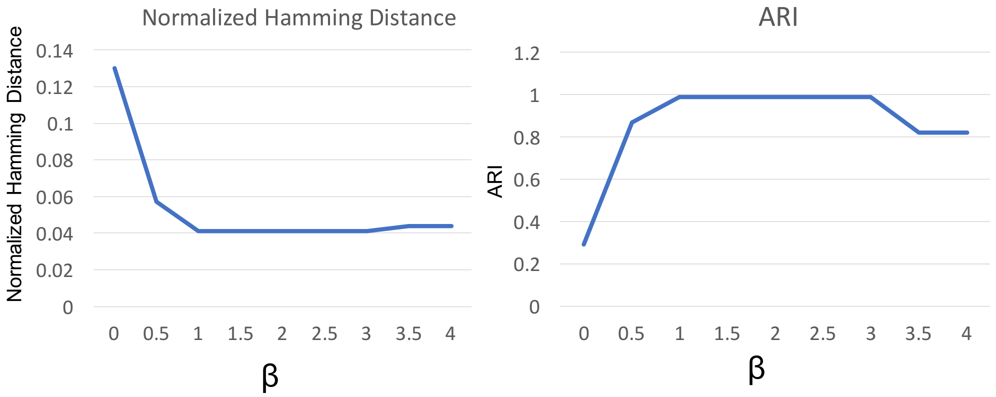


Figure 1: (a) Normalized Hamming distance between true copy number profiles and copy number profiles inferred by STARCH on simulated data for different values of $\beta$. (b) Adjusted Rand Index (ARI) between true clone assignments and the clone assignments inferred by STARCH on simulated data for different values of $\beta$.

the average silhouette score for a range of $K$ and selecting the value of $K$ with the highest average silhouette score.

We then initialize the copy number profile matrix $\mathbf{C}^0$. For each clone $k$ and each bin $j$, we compare the observed expressions in bin $j$ of spots assigned to clone $k$ to the expressions in bin $j$ of normal spots. The distance between these samples is measured by a two-sample one-sided Kolmogorov–Smirnov (KS) test. If the spots in clone $k$ have higher average expression than the normal spots in bin $j$ ($p \leq 10^{-5}$), then we set $c_{kj} =$ Amplification; alternatively if the spots in clone $k$ have significantly lower expression than normal spots ($p \leq 10^{-5}$) then we set $c_{kj} =$ Deletion; otherwise we set $c_{kj} =$ Neutral.

The parameters $\Theta = \cup_{s \in S}\{\mu_s, \sigma_s\}$ are then initialized to the means and standard deviations of bins assigned to each state $s$. The transition probability matrix $T$ is initialized,

$$\begin{bmatrix} 1-2t & t & t \\ t & 1-2t & t \\ t & t & 1-2t \end{bmatrix}$$

where $t = 10^{-5}$. The starting probabilities $\pi$ are initialized to the uniform distribution.

### 7 Runtime

We find that STARCH runs significantly faster than inferCNV on pseudo-spatial transcriptomics data from the breast cancer biopsy S501 (Figure 2).

### 8 Pseudo-spatial transcriptomics data from high grad serous carcinoma cell line OV2295

We generate a synthetic pseudo-spatial transcriptomics dataset from a high grade serous carcinoma cell line, OV2295 obtained from Campbell et al. [2019]. The dataset contains two clones (C and D) as well as their corresponding copy number profiles derived from scDNA-seq. We define $\mathbf{G}$ to be the expression profile

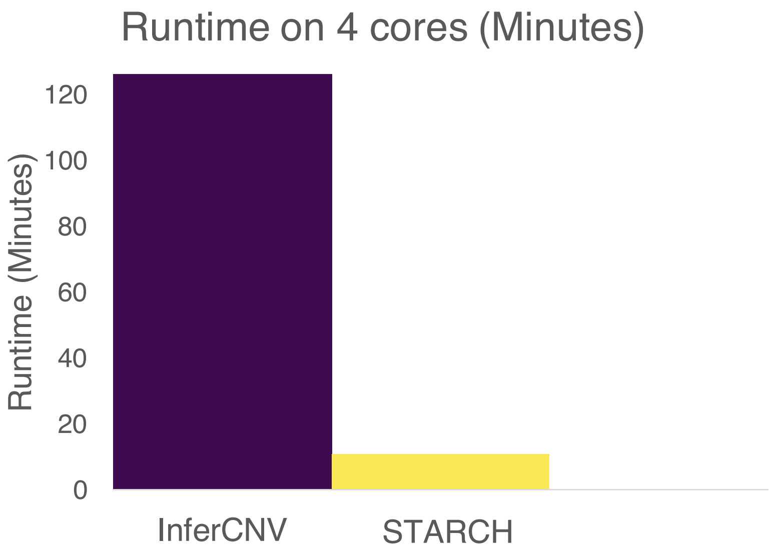


Figure 2: Running time of InferCNV and STARCH on a 4-core Xeon E5-2690 clocked at 2.60GHz using pseudo-spatial data from breast cancer biopsy S501.

derived from scRNA-seq. We use the two clones (C and D) and their corresponding copy number profiles derived from scDNA-seq to simulate the spatial relationships between cells. To simulate spatial relationships between cells we aligned the 1460 cells on a grid with one contiguous region containing cells from clone C and other with cells from clone D (Figure 3). To normalize **G** relative to normal cells with diploid copy number profile we randomly selected 200 of the 674 cells assigned to clone C and defined them as normal cells. We run our standard preprocessing procedure (Section 1) using these selected normal cells, resulting in a normalized and binned matrix **X**.

We ran STARCH, STARCH-nonspatial, and InferCNV on the psuedo-spatial transcriptomics dataset using $K = 2$ clones. We found that by incorporating spatial information STARCH outperformed both STARCH-nonspatial and InferCNV at assigning spots to clones and inferring copy number profiles. STARCH inferred clones with high accuracy (ARI=.99), while InferCNV and STARCH-nonspatial had low accuracy (ARI=.31, ARI= .39 respectively) (Figure 3). In addition, STARCH had the lowest error in its inferred copy number profiles, with a copy number distance of .34 compared to InferCNV with .68 and STARCH-nonspatial with .51.

#### 9 Distinguishing cancer spots from normal spots

STARCH requires the gene expression of normal cells in order to transform the binned input expression to clone copy number profiles. The breast cancer tissue from Ståhl et al. [2016] had a corresponding stained image is labelled to indicate which regions contain cancerous and non-cancerous tissue. However, such information may not be available for other spatial transcriptomics datasets. Thus, we designed a heuristic to distinguish normal cells from tumor cells using spatial transcriptomics data. We observe that by projecting spatial transcriptomics data from Ståhl et al. [2016] onto the first principal component and clustering the loadings into two clusters with K-means (K=2) gave a partition of the spots that closely matched the stained imaging data, with the cluster having the higher average expression corresponding to cancer spots and the cluster having the lower average expression corresponding to non-cancer spots. This indicates that the

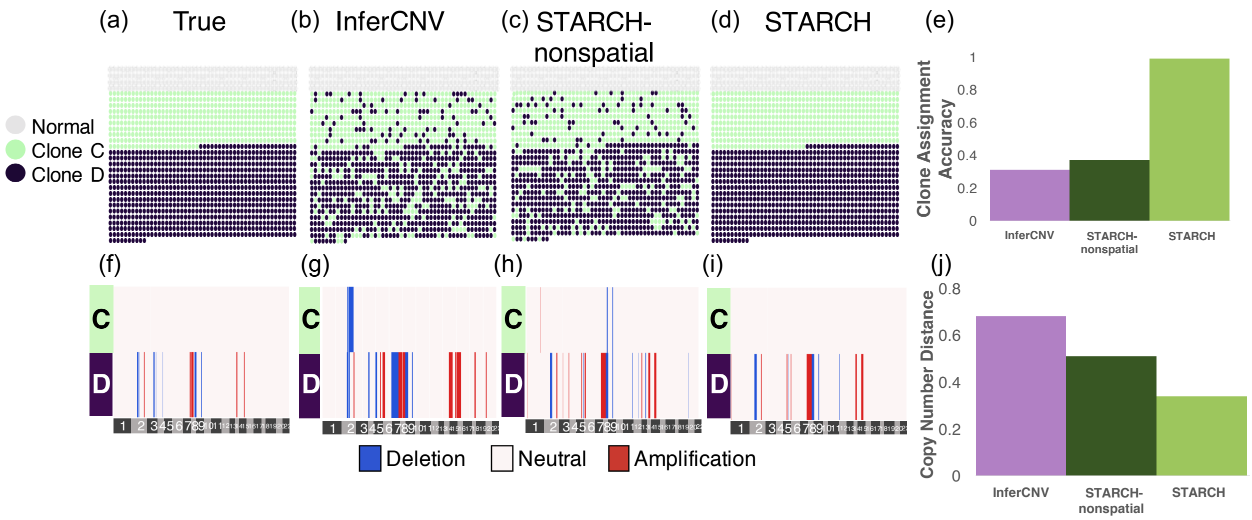


Figure 3: Clone assignment and CNA inference results from InferCNV, STARCH-nonspatial, and STARCH on pseudo-spatial data from high grade serous carcinoma cell line OV2295 [Campbell et al., 2019]. (a-d) Simulated spatial structure with inferred clones C (green), D (purple). Normal spots are indicated in gray. (f-i) Copy number profiles and clones inferred by each method. (e) Clone assignment accuracy for each method measured by Adjusted Rand Index (higher is better). (j) Copy number distance between true and inferred copy number profiles (lower is better).

direction of maximum variance in the expression data corresponds well with the split between normal (non-cancerous) and cancerous spots. We also found that this split was highly correlated with the percent dropout (number of zeros) in each spot, with normal spots having on average higher dropout percent and cancer spots having on average lower dropout percent.

Before performing this heuristic, we preprocessed the expression data as follows:

1. Remove genes with fewer than 15 nonzero entries.
2. For each spot, divide expression of each gene by total gene expression for that spot.
3. Log transform each entry.
4. Z-normalize the gene expression vector for each spot.

We assessed the performance of our heuristic by comparing to the hand-labelled pathologist annotations of the stained imaging data from Ståhl et al. [2016]. We measure both the precision, defined by the percent of correctly predicted normal spots out of all spots predicted normal, and the recall, defined as the percent of normal spots that are recovered out of all the true normal spots. We find that clustering based on the first principal component resulted in high precision ($\geq .9$) at most levels of recall (Figure 4). This performance was consistent across all four layers of the breast tumor biopsy. Clustering based on dropout percent resulted in slightly worse performance, indicating that the first principal component captures some additional variation between the cancer and non-cancer spots. These results motivate us to use the first principal component to distinguish cancer vs. non-cancer spots in spatial transcriptomics data.

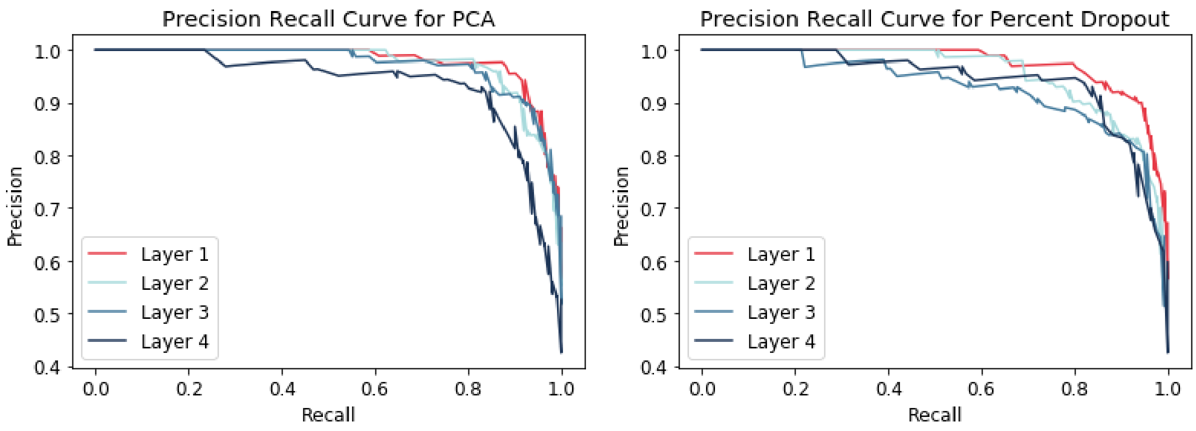


Figure 4: Precision-recall curves for classifying normal spots in spatial transcriptomics data using the first principle component (left) and the percent dropout (right)

#### 10 Negative Control with human heart tissue

As a negative control, we ran STARCH on fresh frozen human heart tissue from a healthy donor obtained from 10X Genomics (`https://support.10xgenomics.com/spatial-gene-expression/datasets/1.0.0/V1_Human_Heart`). The healthy heart tissue should not harbor any copy number aberrations and should contain only normal cells. We ran the full preprocessing pipeline as described in Supplemental Section 1 and applied our procedure for distinguishing tumor from normal spots as described in Supplemental Section 9. Even though the heart tissue should contain only normal spots, our heuristic is guaranteed to identify a cluster of normal and tumor spots, so in this case it will erroneously assign some spots as tumor spots. Despite the classification method misclassifying 2112 of the 4235 spots as tumor spots, when we ran STARCH with $K = 3$, the method output a single clone containing only four small CNAs, with the largest CNA spanning only 4 bins (Figure 5). This low false positive rate demonstrates that even when provided a poor definition of normal and tumor spots, STARCH did not infer many erroneous CNAs. Additionally, even though STARCH was run with $K = 3$ clones, it output a single clone and did not infer erroneous subclones. These two results are promising and suggest that the preprocessing pipeline and the STARCH algorithm will produce few false positive results on real tumor STRNA-seq data.

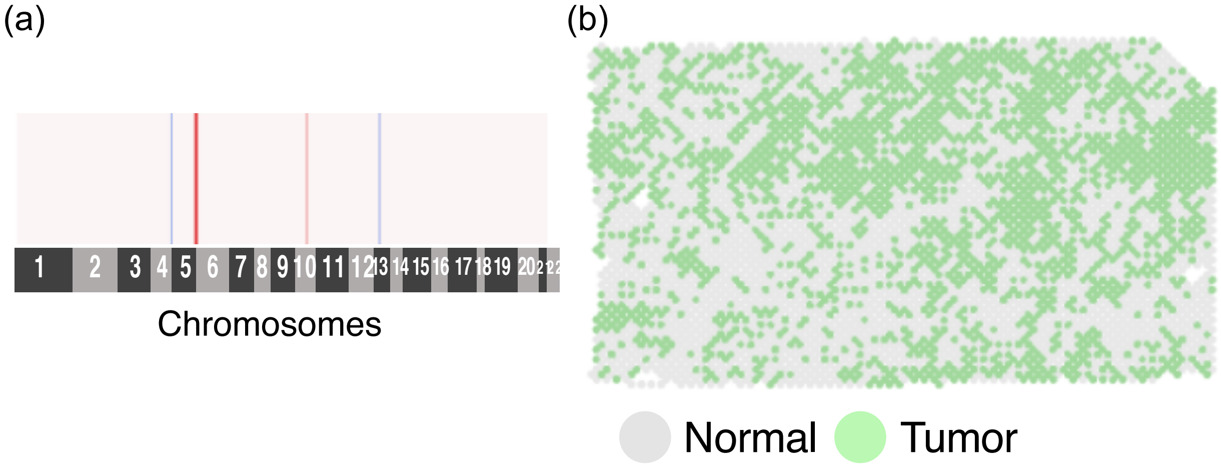


Figure 5: Results of running STARCH on fresh frozen human heart tissue from a healthy donor obtained from 10X Genomics. STARCH was run with $K = 3$ clones but output a single clone. (a) Copy number profile inferred of the tumor spots . (b) Tumor and normal spots visualized on the tissue section.
